# Dispersal evolution diminishes the negative density dependence in dispersal

**DOI:** 10.1101/2020.06.07.138818

**Authors:** Abhishek Mishra, Partha Pratim Chakraborty, Sutirth Dey

## Abstract

In many organisms, dispersal varies with the local population density. Such patterns of density-dependent dispersal (DDD) are expected to shape the dynamics, spatial spread and invasiveness of populations. Despite their ecological importance, empirical evidence for the evolution of DDD patterns remains extremely scarce. This is especially relevant because rapid evolution of dispersal traits has now been empirically confirmed in several taxa. Changes in DDD of dispersing populations could help clarify not only the role of DDD in dispersal evolution, but also the possible pattern of subsequent range expansion. Here, we investigate the relationship between dispersal evolution and DDD using a long-term experimental evolution study on *Drosophila melanogaster*. We compared the DDD patterns of four dispersal-selected populations and their non-selected controls. The control populations showed negative DDD, which was stronger in females than in males. In contrast, the dispersal-selected populations showed density-independent dispersal, where neither males nor females exhibited DDD. We compare our results with previous evolutionary predictions that focused largely on positive DDD, and highlight how the direction of evolutionary change depends on the initial DDD pattern of a population. Finally, we discuss the implications of DDD evolution for spatial ecology and evolution.

## 1. Introduction

Biological dispersal, an integral part of the life history in many taxa (Bonte and Dahirel 2017), is a major determinant of spatial distribution of living organisms. Dispersal patterns influence several ecological phenomena, including biological invasions, range expansions and community assembly (Bowler and Benton 2005; Clobert et al. 2009; Lowe and McPeek 2014). The specifics of a given dispersal event, in turn, are regulated by numerous biotic and abiotic factors (Bowler and Benton 2005; Matthysen 2012). One such factor that can be a prominent cause of variation in the dispersal patterns of many species is the local population density, leading to density-dependent dispersal (DDD) (reviewed in Matthysen 2005; Harman et al. 2020).

Population density can affect the movement of individuals in many ways. For instance, DDD is defined as positive when the per capita dispersal increases with increasing population density, often manifesting as greater proportional movement from dense regions to sparse regions (e.g. Aars and Ims 2000; De Meester and Bonte 2010; Bitume et al. 2013; Lutz et al. 2015). Similarly, negative DDD implies a reduction in per capita movement with increasing population density, resulting in greater aggregation in crowded regions and higher net emigration from sparse regions (e.g. Andreassen and Ims 2001; Baguette et al. 2011; Pennekamp et al. 2014; Mishra et al. 2018b). In addition, recent empirical studies have also reported non-linear DDD, i.e. a combination of positive and negative DDD occurring at different density ranges. Here, U-shaped DDD involves negative DDD at low densities and positive DDD at high densities (e.g. Kim et al. 2009; Fronhofer et al. 2015; Maag et al. 2018), whereas hump-shaped DDD has positive DDD at low density and negative DDD at high density (e.g. Jacob et al. 2016; Chatelain and Mathieu 2017). The classical view has been that positive DDD is more prevalent than the other DDD patterns (Matthysen 2005). This was also supported by a recent literature review on density-dependent emigration, which identified positive DDD as the most common one, followed by negative DDD and non-linear DDD patterns (Harman et al. 2020).

Despite the wealth of experimental studies that have characterized DDD in various taxa, there remains a lack of empirical evidence for the ecological role played by DDD in a spatial context. This is particularly true for the cases where some or the other form of spatial selection, i.e. dispersal evolution at population margins (Phillips et al. 2010; Williams et al. 2019), is involved. Over the past few years, empirical studies across a range of taxa have demonstrated that dispersal traits can evolve rapidly (e.g. Fronhofer et al. 2014; Williams et al. 2016; Weiss-Lehman et al. 2017; Tung et al. 2018b). If rapid dispersal evolution is indeed the norm, it stands to reason that context-dependent features of dispersal, such as DDD, could undergo evolutionary changes too. Furthermore, DDD itself could modulate the spatial selection faced by expanding populations and their subsequent expansion.

In this context, Simmons and Thomas (2004) used wild populations of bush crickets to show that the individuals at the range fronts exhibited a different DDD response (negative DDD) than the individuals at the range core (no DDD). Since then, the general consensus from studies has been that individuals at range edges are expected to evolve less positive or even negative DDD compared with those at range cores (Travis et al. 2009; Fronhofer et al. 2017; Weiss-Lehman et al. 2017). This prediction was empirically confirmed in a study using the protist, *Tetrahymena thermophila* (Fronhofer et al. 2017), where the slope of the density reaction norm changed from slightly positive to neutral or negative. To our knowledge, this is the only empirical study that has demonstrated an evolutionary change in DDD. However, the fact that these results were observed in clonally reproducing *Tetrahymena* strains meant the lack of two things: a) initial standing genetic variation, and b) sexual reproduction, thereby precluding evolution via assortative mating of highly dispersive individuals (i.e. spatial sorting) (Shine et al. 2011). As these two features are central characteristics of dispersing populations in many species, there is a need for empirical investigation of DDD evolution in sexually reproducing populations with standing genetic variation. Moreover, it would allow the investigation of sex-specific changes, if any, during DDD evolution.

Here, we examined the evolutionary changes in DDD using large (breeding population size ~2400 individuals), laboratory-maintained populations of *Drosophila melanogaster*. We used four dispersal-selected populations from a long-running evolutionary experiment, along with their corresponding non-selected control populations. The selected populations underwent dispersal evolution every generation, with 75 generations (~3 years) of selection completed at the time of this study. By comparing the DDD patterns of dispersal-selected and control populations, we could test whether dispersal evolution strengthened or weakened the DDD response. Furthermore, we examined how the sex differences in DDD are affected by evolution. Our results showed a significant negative DDD in the control populations, similar to the ancestral populations. The dispersal-selected populations, however, revealed a complete loss of DDD, suggesting that dispersal evolution had significantly weakened the negative DDD pattern. This was accompanied by a significant upward shift of the global reaction norm across the various densities and a reduction of sex differences in dispersal. We discuss how our results compare to the previous predictions in this context, and the possible implications for dispersal biology.

## 2. Methods

### 2.1 Fly populations and dispersal selection

We used eight laboratory-maintained populations of *D. melanogaster* in this study. Four of these populations (VB_1-4_) had undergone selection for higher dispersal for ~75 generations at the time of the experiment. The other four populations (VBC_1-4_) served as the corresponding controls for the VB populations, i.e. they were reared and maintained under similar conditions for the same number of generations, but not selected for higher dispersal. The subscripts for VB and VBC populations (i.e. 1–4) denote their ancestry, such that populations with the same subscript were derived from the same ancestral population (e.g. VB_1_ and VBC_1_, and so on). The detailed maintenance and selection regime for these populations is available elsewhere (Tung et al. 2018a; Tung et al. 2018b). In brief, all eight populations were maintained at large populations sizes (~2400 individuals) to avoid inbreeding, under a 15-day discrete generation cycle. They had access to 24-h light and experienced a uniform temperature of 25 °C. Every generation, ~50% of the flies (i.e. those that successfully complete dispersal) were selected from the VB populations using replicate two-patch (*source-path-destination*) setups, whereas no such selection was imposed on the corresponding VBC populations (Fig. 1). The length of the path between the *source* and the *destination* was increased intermittently across generations, which mimicked increasing habitat fragmentation over time. As a result of the dispersal selection, VB populations evolved to have a higher dispersal propensity, ability and speed (Tung et al. 2018a; Mishra et al. 2020).

**Fig. 1.**
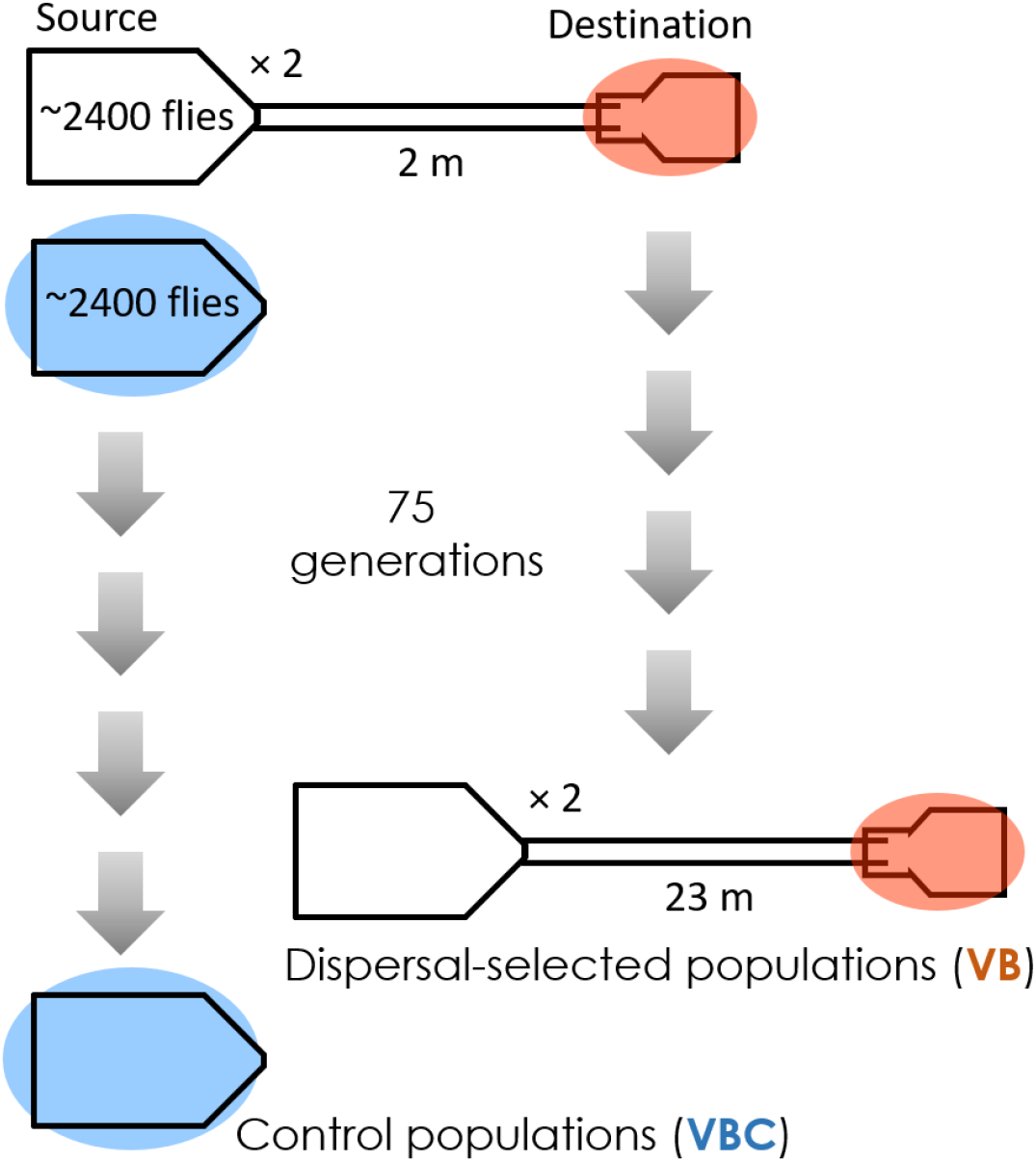
Dispersal selection protocol. Each generation, flies from the four VB (dispersal-selected) populations were subjected to the above dispersal selection protocol. For this, ~2400 adult flies were introduced into the *source* container (1.5 L) of a dispersal setup, which opened into a *path* (plastic tube, ~1 cm diameter). The flies could thus emigrate from the *source* container into the *path*, which then led into the *destination* container (1.5 L). To maintain the population size across generations, two replicate dispersal setups (with ~2400 flies each) were used per population block (e.g. VB1), and the first 50% flies that completed dispersal were selected as the parental population for the next generation. The *path* length was increased intermittently across generations to mimic increasing habitat fragmentation, such that the *path* length was 2 m at generation 0, and 23 m at generation 75. The VBC (non-selected control) populations were maintained identically to the VB populations under the same conditions, but not subjected to the dispersal selection protocol.

### 2.2 Dispersal setup and assay

We used two-patch *source-path-destination* setups to study the dispersal of the flies. This setup was similar to the one used for selection of higher dispersers in VB populations (Tung et al. 2018b), with the only difference being in the size of the containers used as *source* and *destination*. Following an earlier protocol (Mishra et al. 2018b; Mishra et al. 2020), we used 100-mL conical glass flasks as *source* containers and 250-mL plastic bottles as *destination* containers. The source and the destination were connected by a 2-m long *path* (transparent plastic tube; inner diameter ~1 cm) (Fig. 2). We introduced adult flies into the *source* and allowed them to disperse through the *path* into the *destination* for a period of 2 h. We replaced the *destination* container with a fresh, empty container every 15 min during this period, with minimum possible disturbance to the rest of the setup. The number and sex of dispersers during each of these 15-min intervals were manually recorded, to estimate the dispersal propensity and temporal profile of dispersers (see section 2.5 for more details).

**Fig. 2.**
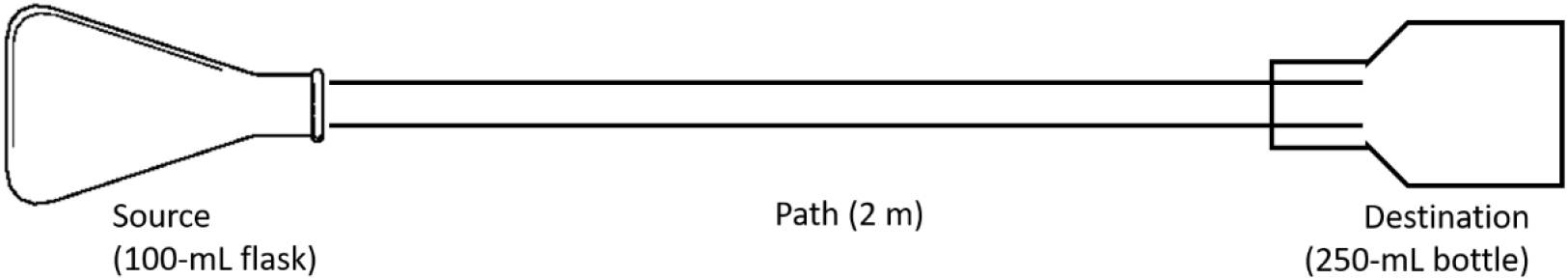
Two-patch dispersal setup used in the study. Adult individuals of *Drosophila melanogaster* were introduced in the *source* container (100-mL conical glass flask), the opening of which was connected to the *path* (2-m long transparent plastic tube of internal diameter ~1 cm). The other end of the *path* opened into the *destination* container (250-mL plastic bottle), with a protrusion of ~3 cm to prevent backflow of successful dispersers into the *path*. During the dispersal assay, the *destination* container could be replaced periodically with a fresh container, to estimate the temporal profile of successful dispersers.

### 2.3 Culturing flies for the experiment

To minimize the contribution of non-genetic parental effects, we reared all VB and VBC populations for one generation under common conditions prior to the experiment. Following an earlier protocol (Mishra et al. 2018b; Mishra et al. 2020), we then generated the flies such that they were age-matched at the time of dispersal assay, and without any apparent confounds of kin-or inbreeding-related effects (Supplementary Text S1). Finally, to eliminate any confounding effects of habitat quality during the dispersal assay, the dispersal setup (Section 2.2) was devoid of any food or moisture (similar to Mishra et al. 2018b; Mishra et al. 2020).

### 2.4 Experimental design

For comparing the pattern of DDD between VB (dispersal-selected) and VBC (control) populations, we assayed each of the four VB populations with its corresponding control population (i.e. VB_1_ assayed with VBC_1_, and so on). Four dispersal density treatments were used, namely 60, 120, 240 and 480 individuals per *source* container. These densities were chosen based on an earlier DDD study on the ancestral populations of these flies (Mishra et al. 2018b). As density can affect the physiology and dispersal of individuals in multiple possible ways and across life stages (Matthysen 2005; Harman et al. 2020), we decided to limit our study to adult density and adult dispersal. With a uniform rearing until the adult stage (Supplementary Text S1), this ensured that adult density was the only environmental factor that differed among these four treatments.

We performed the experiment over multiple consecutive days, with a fresh set of age-matched (12-day-old from egg-lay) adult flies every day (Supplementary Text S1). This way, all the density treatments for both VB and VBC populations could be assayed together every single day, allowing a complete replication of all the treatments each day. The four population blocks (i.e. 1–4) were assayed one after the other, wherein each block was assayed over 4 consecutive days. As a result, the entire experiment ran over the course of 16 days (4 blocks × 4 days, allowing us to obtain a total of 16 replicates (blocked by population block and day) for each density treatment of VB and VBC.

As mentioned above, 12-day-old (from egg-lay) adult flies were used for the dispersal assay. On day 11 of their age (21 h prior to the dispersal assay), the flies from the relevant populations were separated by sex under light CO_2_ anesthesia, and then randomly assigned to the four density treatments (60, 120, 240 and 480) with a strict 1:1 sex ratio. Thus, the density treatment of 60 individuals comprised 30 males and 30 females, and so on. The prepared sets of flies were then transferred into separate 100-mL conical glass flasks (identical to the source in the dispersal setup) containing ~35 mL of banana-jaggery medium. The next day (at 12^th^ day of age), these flies were assayed for their dispersal as described in Section 2.2 (Fig. 2). As we assayed one replicate of each density for VB and VBC populations per day, a total of 28,800 flies (2 populations × 4 blocks/population × 4 days/block × 900 flies/day) were used for this experiment.

### 2.5 Dispersal traits

For every dispersal setup (replicate), we counted the number of male and female flies that reached the destination during each of the 15-min intervals until the end of dispersal assay (2 h). Moreover, we also recorded the number and sex of flies that emigrated from the *source* but did not reach the *destination*, i.e. those found within the *path* tube at the end of the dispersal assay.

These data allowed us to estimate the dispersal propensity, i.e. the proportion of flies that initiated dispersal from the *source* (Friedenberg 2003), as:

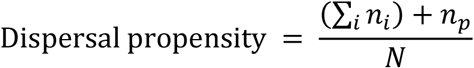

where *n_i_* is the number of flies that reached *destination* during the *i*^th^ 15-min interval, *n_p_* is the number of flies found in the *path* at the end of dispersal assay and *N* is the total number of flies introduced in the setup (i.e. 60, 120, 240 or 480).

In addition, we obtained the overall temporal profile of dispersers for each replicate. For this, we calculated the proportion of dispersers that reached *destination* during each 15-min interval, relative to the final number of successful dispersers (Mishra et al. 2018b; Mishra et al. 2020).

### 2.6 Statistical analyses

As stated in section 2.4, the experiment involved four blocks of VB and VBC populations, each of which was assayed over 4 consecutive days, yielding 16 replicates in total for each density treatment. As one replicate for each of the four density treatments was assayed per day, *day* (1-4) was included as a random factor that was nested inside *block* (1-4), another random factor, for all analyses. This was done to account for any day-to-day microenvironmental variations.

The dispersal propensity data were first analyzed together using a Generalized Linear Mixed Model (GLMM) with binomial error distribution (and logit link function) for the dispersal propensity data, using the ‘glmmPQL’ function from the ‘MASS’ package v7.3-51.4 (Ripley et al. 2013) in R v3.6.2 (R Core Team 2019). In addition to the random factors *day* and *block* mentioned above, the model incorporated *dispersal selection* (VB and VBC), *density* (60, 120, 240 and 480) and *sex* (male and female) as fixed factors. Following this, the pairwise differences of interest were assessed using individual GLMMs. Here, *sex* (male/female) was used as the fixed factor for determining sex-biased dispersal at a given density treatment for a population (e.g., at density 60 of VBC population). Similarly, *density* was used as the fixed factor for pairwise comparisons of density treatments of a given sex in a population, to ascertain the direction of its DDD pattern (e.g. comparison between densities 60 and 120 for VBC females). The familywise error rate for these pairwise GLMMs was then corrected using the Holm-Šidák correction (Abdi 2010). Cohen’s *d* was used as a measure of effect size or pairwise differences, with the effect interpreted as large, medium and small for *d* ≥ 0.8, 0.8 > *d* ≥ 0.5 and *d* < 0.5, respectively (Cohen 1988).

For analyzing the temporal profile data of VB and VBC dispersers of each sex, we carried out similar GLMMs in R, but this time with *density* (60, 120, 240 and 480) and *time* (15, 30, 45, 60, 75, 90, 105, and 120 min) as fixed factors. The random factors, i.e. *day* nested within *block*, remained the same as above.

## 3. Results

The GLMM for dispersal propensity data yielded a significant effect of *selection × density* (t = −5.30, p < 10^−4^) as well as *selection × density × sex* (t = 2.98, p = 0.003) (Supplementary Table S1). We thus limit our interpretation to the final three-way interaction term. Following this, we used pairwise GLMMs with familywise-error corrections to assess the DDD and instances of sex-biased dispersal in these populations (Section 2.6). We report the interpretations of these analyses here, with the detailed results presented in Supplementary Tables S2–S4.

### 3.1 Negative and sex-biased DDD observed in control populations

VBC populations showed a negative DDD, where the dispersal propensity was lower at higher densities (Fig. 3A). Furthermore, the strength of negative DDD was stronger in VBC females than in VBC males (Fig. 3A, Supplementary Table S2). As a result, a significant female-biased dispersal was observed at densities 60 and 120, whereas a significant male-biased dispersal was observed at the highest density, 480 (Fig. 3A, Supplementary Table S3). Taken together, these results indicate a strong negative DDD in VBC populations, along with a sex-biased asymmetry: females showed a stronger negative DDD response than the males.

**Fig. 3.**
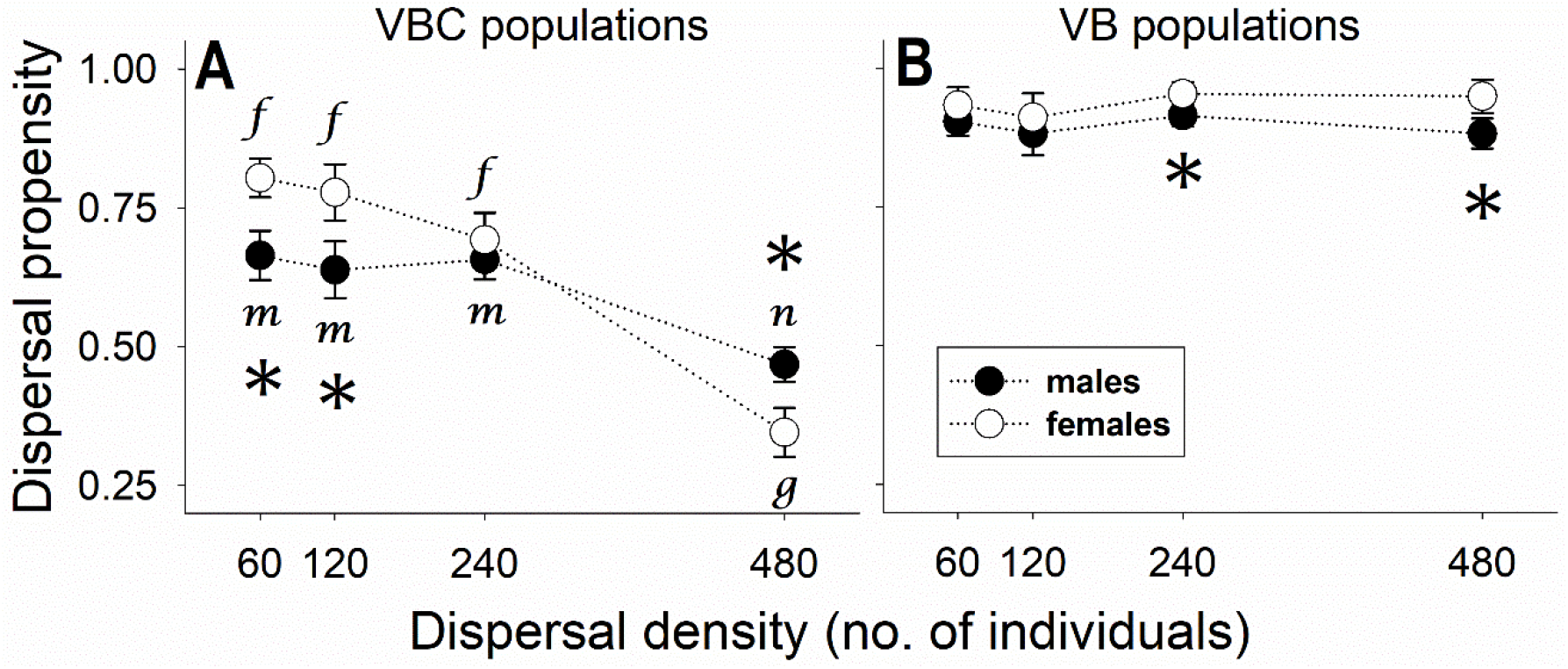
Effect of dispersal evolution on density-dependent dispersal (DDD). Mean dispersal propensity (± SE) across the four density treatments for (A) VBC (control) and (B) VB (dispersal-selected) individuals. A strong negative DDD was observed for VBC individuals, but the VB individuals showed no DDD. Open circles denote the data for females, with lower-case letters starting from *f* representing the significant differences in female dispersal propensity across densities. Closed circles denote the data for males, with lower-case letters starting from *m* representing the significant differences in male dispersal propensity across densities. Asterisks (*) denote significant sex-biased dispersal (p < 0.05) at a given density. See Supplementary Tables S2 and S3 for the exact p values and the associated effect sizes.

The effect of density for VBC populations was apparent in the temporal dispersal profile as well. We found a significant effect of *density* on the female dispersers (Fig. 4B, Supplementary Table S4.2), while the male dispersal data showed a significant effect of *density* as well *density × time* interaction (Fig. 4A, Supplementary Table S4.1). In both sexes, a majority of dispersal (> 50%) occurred within one hour for the first three density treatments (i.e. 60, 120, and 240 individuals), whereas a majority of dispersal took place after the first hour in the highest density treatment (480 individuals) (Figs. 4A and 4B). Taken together, this means that the dispersal propensity of VBC individuals was not only suppressed at higher density (Fig. 3A), but the dispersers also took longer to complete their journey from the *source* to the *destination* (Figs. 4A and 4B).

**Fig. 4.**
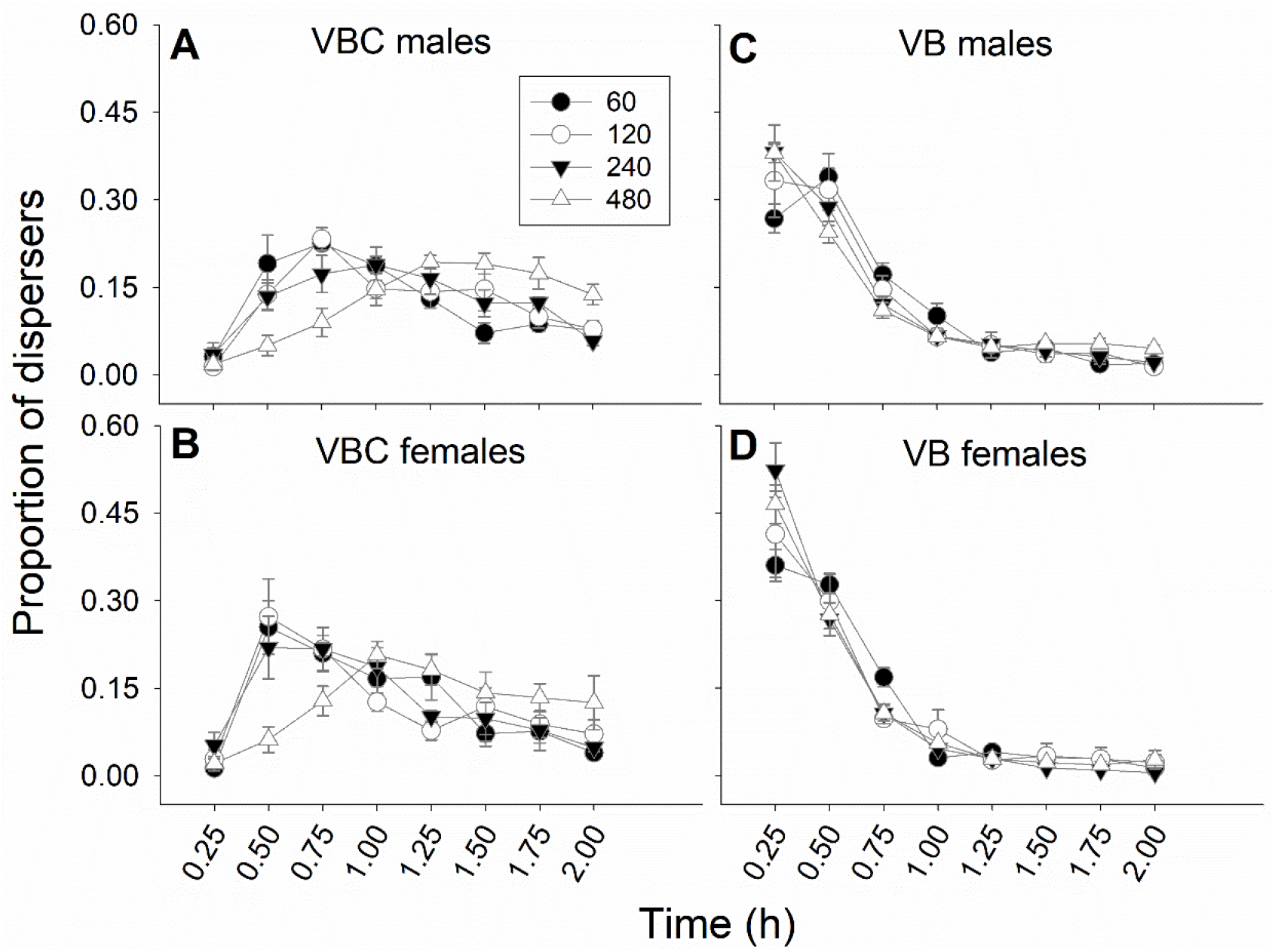
Temporal profile of VB (dispersal-selected) and VBC (control) flies across the four density treatments. Average proportion (± SE) of individuals completing dispersal during the specified temporal bins for (A) VBC males, (B) VBC females, (C) VB males and (D) VBC females. As can be observed, both male and female dispersers of VB populations completed the majority (> 50%) of dispersal within the first two temporal bins (i.e. by 0.5 h). In contrast, the VBC dispersers had a more staggered dispersal, thereby taking longer to complete 50% dispersal. See Supplementary Table S4 for the exact p values.

### 3.2 No DDD observed for either sex in dispersal-selected populations

In contrast to the results for VBC, we found that the VB populations did not show a DDD response at all. Neither males nor females showed a significant difference in their dispersal across the four treatment densities (Fig. 3B, Supplementary Table S1). Moreover, the only instances of sex-biased dispersal were that of female-biased dispersal at the two highest densities (i.e. 240 and 480) (Fig. 3B, Supplementary Table S2), in contrast to the sex-unbiased and male-biased dispersal observed at these densities in VBC populations, respectively (*cf.* Fig. 3A and 3B).

In terms of the temporal dispersal profile as well, VB flies in all density treatments completed the dispersal much faster than the VBC flies (*cf.* Figs. 4A, 4B with Figs. 4C, 4D). As a result, within the first 30 minutes of the dispersal run, a majority of VB dispersers (> 50%) had completed their journey from the *source* to the *destination* (Figs. 4C and 4D). Furthermore, neither male nor female dispersal had a significant effect of *density* or *density × time* interaction (Supplementary Tables S4.3 and S4.4). Taken together, the results suggest that both dispersal propensity and the speed of dispersal of VB populations were largely unaffected by the density treatments.

## 4. Discussion

### 4.1 Negative DDD in control populations

The non-selected control (VBC) populations showed a significant negative DDD, as the dispersal propensity showed a decreasing trend with increasing density (Fig. 3A). Furthermore, this response was significantly stronger in females than in males. Because of this asymmetry between the sexes, there was a switch in the direction of sex-biased dispersal: we observed a significant female-biased dispersal at low densities but a significant male-biased dispersal at high densities (Fig. 3A). Finally, the significant reduction in dispersal propensity was also accompanied by considerably delayed dispersal at the highest density (i.e. 480 individuals) (Figs. 4A and 4B). All these results are in line with those observed for the ancestral populations of VBC and VB flies, assayed under similar conditions (Mishra et al. 2018b). This consistency of results for ancestral and VBC populations confirms that the DDD response of these flies has remained relatively unchanged over the course of separate rearing for 75 generations (~3 years).

The DDD response of the dispersal-selected (VB) populations would thus be compared against this expectation of a negative DDD pattern. While empirical studies on the evolution of DDD are extremely rare (although see Fronhofer et al. 2017), the general prediction from studies is that populations undergoing rapid dispersal evolution should evolve higher dispersal rates at low densities. This is predicted to cause an abatement of positive DDD patterns and a likely emergence of negative DDD (Travis et al. 2009; Fronhofer et al. 2017; Weiss-Lehman et al. 2017). Since negative DDD was already seen in VBC populations, the next step was to see whether dispersal evolution in VB populations had indeed strengthened this negative DDD, as per the expectation from literature.

### 4.2 Absence of DDD in dispersal-selected populations

In contrast to the negative DDD observed in VBC populations, we observed a complete absence of DDD in VB populations. Both sexes in VB populations showed a consistently high dispersal propensity of >80% across the four densities (Fig. 3B). While there was an increase in dispersal propensity at low densities, as predicted by earlier studies (Travis et al. 2009; Fronhofer et al. 2017), it was also accompanied by a much larger increase in the dispersal propensity at high densities (Fig. 3B). As a result, we obtained an almost flat DDD response, where a consistently high dispersal occurred irrespective of the density treatments.

In addition to the loss of negative DDD, the VB populations also narrowed the extent of sex differences in dispersal (*cf.* Figs. 3A and 3B). Sex-biased dispersal in VB populations was observed only at the highest two densities, where a significant female-biased dispersal was observed. This is in direct contrast to the male-biased dispersal observed for VBC populations at the highest density (i.e. 480) (*cf.* Figs. 3A and 3B). While the evolution of sex-biased dispersal remains a prominent topic of investigation, to our knowledge, this is the first empirical demonstration of an evolutionary change in sex-biased dispersal, that too under identical environmental conditions.

So what explains the change in DDD by dispersal evolution and the loss of sex differences in VB populations? The VB populations have undergone continuous selection for higher dispersal via spatial sorting every generation. The selection setup is such that females need to complete the *source*-to-*destination* dispersal in order to contribute to the next generation, whereas males could theoretically mate with several females before the dispersal run and still be able to sire progeny in the next generation (Fig. 1; Tung et al. 2018b)). The high selection pressure on dispersal speed (only the first 50% dispersers are selected each generation) means that, over generations, the rate at which the eventually successful female dispersers would leave the *source* container would tend to increase. Additionally, leaving the source container quickly can also allow the females to escape excessive harassment by males (e.g. Byrne et al. 2008; Yun et al. 2017; Malek and Long 2019). However, the high female dispersal thus evolved could create a shortage of mates for males, likely resulting in increasingly higher mate-finding dispersal by them (Shaw and Kokko 2014; Mishra et al. 2020). This way, spatial sorting would then ensure that both males and females evolve increasingly similar dispersal kernels (Meier et al. 2011; Shaw and Kokko 2014) with high dispersal propensity, ability and speed. As a result, the dispersal-selected females can emigrate away to escape the excessive mate harassment experienced at high densities, while the males track their movement via mate-finding dispersal (Fig. 3B, Figs. 4C and 4D). Over generations, the continuous evolution of higher dispersal could thus lead to a similar magnitude of DDD loss in both sexes.

### 4.3 Implications

Our results showed that dispersal evolution completely reversed the negative DDD seen in control and ancestral populations. This was accompanied by a loss of sex differences in dispersal, leading to fewer instances of sex-biased dispersal. Below, we discuss some ecological and evolutionary implications of these results.

First, our results demonstrate a strong effect of dispersal evolution, resulting in uniformly high dispersal of males and females irrespective of the environmental context. In other words, this is yet another piece of strong evidence that dispersal evolution leads to context-independent dispersal (Tung et al. 2018b). As a result, we predict that populations undergoing strong spatial selection would exhibit increasingly more phenotype-dependent movement than context-dependent movement (*sensu* Clobert et al. 2009; Clobert et al. 2012).

Second, the narrowing of sex differences confirms that males and females can evolve strikingly similar dispersal patterns, even if their initial dispersal patterns are very different. In species where dispersal costs differ significantly between males and females (Gros et al. 2008; Trochet et al. 2016; Mishra et al. 2018a), this could alter the physiology and life history of either or both sexes.

Third, the evolutionary change in DDD observed here, i.e. DDD becoming less negative, at first seems contrary to the earlier predictions (Travis et al. 2009; Fronhofer et al. 2017; Weiss-Lehman et al. 2017). However, a closer examination of the issue reveals some interesting points about DDD evolution. For instance, we note that the direction of this evolutionary change (i.e. increase or decrease in the magnitude of positive or negative DDD) is likely dependent on the shape of initial DDD. The populations modeled by Travis et al. (2009) had positive DDD initially, which then evolved density-independent dispersal with time. Similarly, the *Tetrahymena* populations used by Fronhofer et al. (2017) showed a slight positive DDD, which turned neutral or negative upon evolution. In our case, evolution resulted in the abatement of a strong negative DDD pattern (Fig. 3). A common theme, therefore, is that evolution can counter the prevalent mode of DDD, thereby making dispersal less density-dependent. As a result, the question for future studies would likely be whether evolution enforces or diminishes the initial DDD pattern, rather than the absolute direction of change (i.e. towards or away from positive or negative DDD).

Finally, DDD may or may not interact with other phenomena affected by density, such as density-dependent growth and selection. In the case of Fronhofer et al. (2017), dispersal was especially beneficial at low densities, and this differential selection resulted in a negative DDD. In contrast, the selection pressure on dispersal in our case was largely density-independent (Fig. 1). We speculate these interactions to further amplify when the density effects in a population are even more prominent. For instance, the ‘pulled’ vs. ‘pushed’ wave models of invasion invoke the absence and presence of Allee effects at range edges, respectively (Roques et al. 2012; Andrade-Restrepo et al. 2019). As these invasion waveforms can themselves undergo evolutionary transitions (Gandhi et al. 2016; Erm and Phillips 2020), the underlying DDD patterns are expected to shift rapidly as well. In fact, DDD has been recently proposed as a putative mechanism for these transitions (Dahirel et al. 2020), highlighting yet another important way in which DDD and dispersal evolution interact with each other.

## Acknowledgements

AM was supported by a Senior Research Fellowship from the Council of Scientific and Industrial Research, Government of India. This study was supported by a research grant (#CRG/2018/001333) from Science and Engineering Research Board, DST, Government of India, and internal funding from IISER-Pune.

## Supplementary Online Material

## Supplementary Text S1: Detailed fly culturing methods

For a given population block (e.g. VB_1_ and VBC_1_), two cages each for the VB and VBC populations were prepared with ~2400 adult flies per cage. The parental generation of these flies had been reared under common conditions for one generation, to minimize the contribution of non-genetic parental effects (Section 2.3 in main text). To collect eggs for a particular day of the assay, one of the two cages for a given population (e.g. VB_1_) was supplied with live yeast plate for ~24 h. Fresh banana-jaggery medium was then supplied to the flies, allowing the females to oviposit for 12 h. After this period, ~50 eggs each were collected into forty 35-mL plastic vials containing ~6 mL of banana-jaggery medium. Following this, a plate with live yeast paste was provided to the flies again, to enable another round of egg collection. The same procedure was followed for the second cage of the given population (VB_1_, from the current example) as well, with a constant difference of 24 h between the cages. Therefore, it was possible to collect eggs, alternatively from two cages of the same population, for 4 days. Following this protocol for the VB and VBC populations of a given population block together, we ensured that we were able to get four independent replicates of VB and VBC flies per population block (Section 2.4 in main text). The large breeding population size (~2400) in each cage ensured that the flies were not inbred, and random sampling from a large number of eggs during their collection reduced the chances of kin being sampled together. Moreover, as flies from a single set of collected eggs were used for the dispersal assay on a particular day, they were all of the same age (12^th^ day from egg collection) at the time of their assay.

---

**Table S1.**
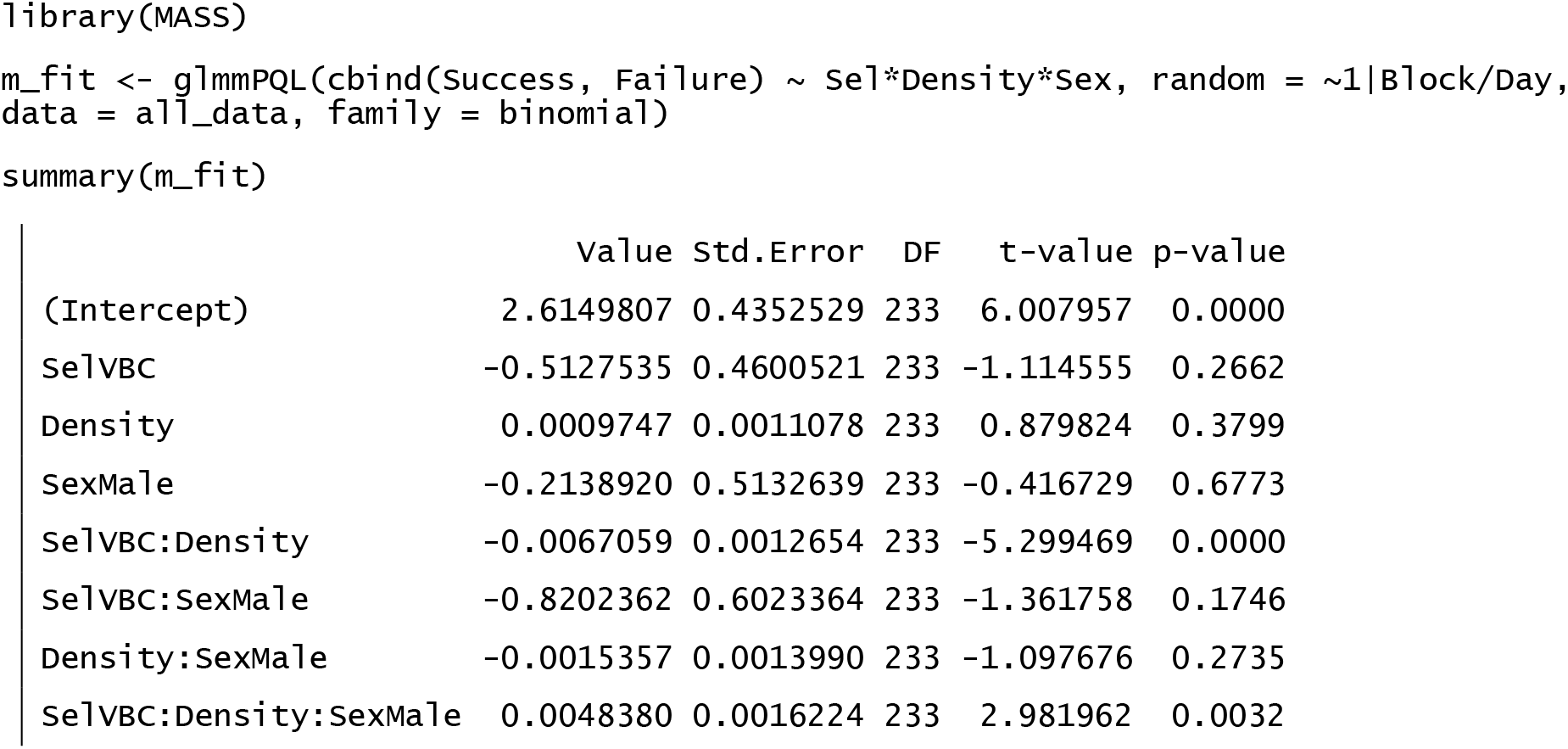
Binomial GLMM results for the dispersal propensity data.

**Table S2.** Within-sex pairwise differences for dispersal propensity across the density treatments for VBC (control) and VB (dispersal-selected populations), after familywise error correction using sequential Holm-Šidák method. For the significant differences (*p* ≤ 0.05), pairwise effect sizes (Cohen’s *d*) are computed.

| Population | Sex | Densities compared | p value | Cohen's $d$ | Effect size interpretation |
| --- | --- | --- | --- | --- | --- |
| VBC<br>(Control) | Males | 60 – 120 | 0.92 | - | - |
|  |  | 60 – 240 | 0.907 | - | - |
|  |  | 60 – 480 | 0.018 | 1.209 | Large |
|  |  | 120 – 240 | 0.97 | - | - |
|  |  | 120 – 480 | 0.014 | 0.954 | Large |
|  |  | 240 – 480 | 0.0006 | 1.316 | Large |
|  | Females | 60 – 120 | 0.65 | - | - |
|  |  | 60 – 240 | 0.22 | - | - |
| | | 60 – 480 | $< 10^{-4}$ | 2.699 | Large |
|  |  | 120 – 240 | 0.22 | - | - |
| | | 120 – 480 | $< 10^{-4}$ | 2.136 | Large |
| | | 240 – 480 | $< 10^{-4}$ | 1.760 | Large |
| VB<br>(Dispersal-selected) | Males | 60 – 120 | 0.82 | - | - |
|  |  | 60 – 240 | 0.92 | - | - |
|  |  | 60 – 480 | 0.95 | - | - |
|  |  | 120 – 240 | 0.86 | - | - |
| | | 120 – 480 | $> 0.99$ | - | - |
|  |  | 240 – 480 | 0.86 | - | - |
|  | Females | 60 – 120 | 0.83 | - | - |
|  |  | 60 – 240 | 0.72 | - | - |
|  |  | 60 – 480 | 0.77 | - | - |
|  |  | 120 – 240 | 0.44 | - | - |
|  |  | 120 – 480 | 0.70 | - | - |
|  |  | 240 – 480 | 0.83 | - | - |

**Table S3.** Sex-bias in dispersal propensity for VBC (control) and VB (dispersal-selected populations). For significant pairwise differences after familywise error correction using sequential Holm-Šidák method (*p* ≤ 0.05), effect sizes (Cohen’s *d*) are computed. m: males, f: females

| Pre-dispersal density | Densities | p value | Cohen's $d$ | Effect size interpretation |
| --- | --- | --- | --- | --- |
| **VBC**<br>**(Control)** | 60m – 60f | $1.2 \times 10^{-3}$ | 0.906 | Large |
| | 120m – 120f | $< 10^{-4}$ | 0.740 | Medium |
|  | 240m – 240f | 0.32 | - | - |
| | 480m – 480f | $1.4 \times 10^{-3}$ | 0.799 | Medium |
| **VB**<br>**(Dispersal-selected)** | 60m – 60f | 0.17 | - | - |
|  | 120m – 120f | 0.40* | - | - |
| | 240m – 240f | $9.9 \times 10^{-3}$ | 0.643 | Medium |
| | 480m – 480f | $8.0 \times 10^{-4}$ | 0.901 | Large |

\*The model for this comparison yielded a singular convergence error. Hence, the nested random factor *day* was dropped from the model for this comparison.

**Table S4.** Binomial GLMM results for the temporal dispersal profile data.

|  | value | Std.Error | DF | t-value | p-value |
| --- | --- | --- | --- | --- | --- |
| (Intercept) | -2.1014609 | 0.23310889 | 493 | -9.014933 | 0.0000 |
| Density | -0.0037685 | 0.00051631 | 493 | -7.298930 | 0.0000 |
| Time | 0.0029012 | 0.00231313 | 493 | 1.254225 | 0.2104 |
| Density:Time | 0.0000217 | 0.00000668 | 493 | 3.243732 | 0.0013 |

|  | value | Std.Error | DF | t-value | p-value |
| --- | --- | --- | --- | --- | --- |
| (Intercept) | -1.3541255 | 0.3635463 | 493 | -3.724768 | 0.0002 |
| Density | -0.0045926 | 0.0005703 | 493 | -8.053237 | 0.0000 |
| Time | -0.0018450 | 0.0028226 | 493 | -0.653644 | 0.5136 |
| Density:Time | 0.0000125 | 0.0000083 | 493 | 1.504847 | 0.1330 |

|  | value | Std.Error | DF | t-value | p-value |
| --- | --- | --- | --- | --- | --- |
| (Intercept) | -0.3225315 | 0.16654551 | 491 | -1.936597 | 0.0534 |
| Density | -0.0005142 | 0.00034961 | 491 | -1.470838 | 0.1420 |
| Time | -0.0129992 | 0.00273115 | 491 | -4.759594 | 0.0000 |
| Density:Time | 0.0000016 | 0.00000712 | 491 | 0.223809 | 0.8230 |

|  | value | Std.Error | DF | t-value | p-value |
| --- | --- | --- | --- | --- | --- |
| (Intercept) | 0.15567018 | 0.21717952 | 459 | 0.716781 | 0.4739 |
| Density | 0.00002975 | 0.00039297 | 459 | 0.075709 | 0.9397 |
| Time | -0.02057236 | 0.00352219 | 459 | -5.840792 | 0.0000 |
| Density:Time | 0.00001496 | 0.00000951 | 459 | 1.572758 | 0.1165 |

